# DNA methylation machinery is involved in development and reproduction in the viviparous pea aphid (*Acyrthosiphon pisum*)

**DOI:** 10.1101/2024.02.14.579807

**Authors:** Kane Yoon, Stephanie Williams, Elizabeth J. Duncan

## Abstract

The pea aphid (*Acyrthosiphon pisum*) like the majority of extant aphids displays cyclical parthenogenesis - the ability of mothers to switch the reproductive mode of their offspring from reproducing parthenogenetically to sexually in response to environmental cues. The pea aphid genome encodes two paralogs of the *de novo* DNA methyltransferase gene, *dnmt3a* and *dnmt3x*. Here we show, using phylogenetic analysis, that this gene duplication event occurred at least 106 million years ago, likely after the divergence of the lineage leading to the Aphidomorpha (aphids, phylloxera and adelgids) from that leading to the scale insects (Coccoidea) and that the two paralogs are maintained in the genomes of all aphids examined. We also show that the mRNA of both *dnmt3* paralogs are maternally expressed in the viviparous aphid ovary. During development both paralogs are expressed in the germ cells of embryos beginning at stage 5 and persisting throughout development. Chemical inhibition of the DNA methylation machinery leads to defects of oocytes and early stage embryos, and causes a proportion of later stage embryos to be born dead or die soon after birth. These phenotypes suggest a role for DNA methyltransferases in reproduction, consistent with that seen in other insects. Taking the vast evolutionary history of the *dnmt3* paralogs, and the localization of their mRNAs in the ovary, we suggest there is a role for *dnmt3a* and/or *dnmt3x* in early development, and a role for DNA methylation machinery in reproduction and development of the viviparous pea aphid.

## Introduction

Pea aphids (*Acyrthosiphon pisum*) are hemimetabolous insects that are voracious pests. Like many species of aphid, they are responsible for huge economic loss through virus-mediated and direct damage to crop plants (Brault et al. 2010; Guerrieri and Digilio 2008). This is due, at least in part, to their ability to reproduce via viviparous parthenogenesis, which allows them to quickly amass large populations of genetically identical, live-birthed daughters (Dixon 1985). While pea aphids and most extant aphids reproduce parthenogenetically in the spring and summer, they parthenogenetically birth oviparous sexual females and males in the autumn and winter triggered as days become shorter and colder (Moran 1992; Simon et al. 2002). This life-history strategy is known as cyclical parthenogenesis, and is a transgenerational polyphenism (Ogawa and Miura 2014). These sexual pea aphids mate and produce overwintering eggs which hatch post diapause as viviparous parthenogenetically reproducing daughters in the spring, and the cycle continues (Dixon 2012; Ogawa and Miura 2014). Sexual reproduction has been totally lost in approximately 3% of aphid species (Dixon 2012) and 30-40% of populations of aphid species show variation in reproductive mode which includes obligate parthenogenesis (Defendini et al. 2023; Jaquiery et al. 2014; Dixon 2012; Moran 1992). Cyclical parthenogenesis is the ancestral reproductive mode of the Aphidomorpha (aphids, phylloxera and adelgids) which evolved approximately 298 million years ago (Grimaldi and Engel 2005) from an ancestor that was oviparous (Gavrilov-Zimin 2021) and sexually reproducing (Dixon 2012). True aphids (Aphididae) subsequently evolved viviparity (Davis 2012) (development occurring inside the mother) between 65-106 million years ago (Johnson et al. 2018). Cyclical parthenogenesis and vivparity are both evolutionary innovations in the aphid lineage. But the mechanisms which control the plasticity of reproductive mode and ovary development in the offspring are largely unknown. Understanding the genes and molecular processes that underpin these innovations is key to our understanding of how this novelty evolves and may also yield unique insight into novel pest control strategies.

Epigenetic mechanisms, heritable and environmentally responsive changes to a genome that are not direct changes to DNA sequence, have been proposed as possible elements mediating plastic responses (Duncan et al. 2022; Richard et al. 2021; Richard et al. 2019). DNA methylation is one such mechanism which involves the reversible addition of methyl groups to nucleotides, primarily to cytosines in CpG dinucleotides (Bewick et al. 2017; Robertson 2005). The functions of DNA methylation are well understood in mammals where CpG methylation is typically constrained to regulatory regions of genes and downregulates gene expression (Li and Zhang 2014). Conversely, relatively little is known about the role of DNA methylation in insects, particularly hemimetabolous insects (Provataris et al. 2018; Bewick et al. 2017; Duncan et al. 2022; Dixon and Matz 2022). Insect DNA methylation occurs primarily in gene bodies and is associated more with highly conserved and stably expressed genes (Bewick et al. 2017; Lewis et al. 2020; Provataris et al. 2018; Sarda et al. 2012). Several studies have experimentally explored possible roles for DNA methylation in insect phenotypic plasticity, yet, a clear general role has not been established (reviewed in Duncan et al. 2022). Intriguingly, several recent studies have strongly suggested methylation independent roles in reproduction for *dnmt1* in range of insects (Blattodea (Ventos-Alfonso et al. 2020), Hemiptera (Shelby et al. 2023; Amukamara et al. 2020; Bewick et al. 2019; Cunningham et al. 2024; Washington et al. 2021), Hymenoptera (Arsala et al. 2022; Ivasyk et al. 2023; Zwier et al. 2012), Lepidoptera (Li et al. 2019a), Coleoptera (Schulz et al. 2018; Gegner et al. 2020)), but it is unclear at this time whether similar non-methylation functions might exist for *Dnmt3*.

The pea aphid has a complete suite of genes encoding the DNA methylation machinery, including one copy of *DNA methyltransferase 1* (*dnmt1*), two paralogs of the *de novo* DNA methyltransferase, *DNA methyltransferase 3* (*dnmt3*): *dnmt3a* and *dnmt3x* (Walsh et al. 2010) and a single copy of the putative demethylation enzyme *tet methylcytosine dioxygenase 1* (*tet1*, Duncan et al. 2022). In the case of the pea aphid *dnmt3* paralogs, *dnmt3a* has been suggested to be a functional as a *de novo* DNA methyltransferase, while *dnmt3x* appears to have diversified, and is lacking some of the key domains thought to be necessary for carrying out DNA methylation (Walsh et al. 2010). This raises the possibility that the duplication of *dnmt3* that gave rise to *dnmt3a* and *dnmt3x* allowed for subfunctionalisation or neofunctionalization (of *dnmt3x*) and a possible role in the novel reproductive and developmental strategy of cyclical viviparous parthenogenetic reproduction.

Here, we demonstrate that the *dnmt3* gene duplication occurred at least 106 million years ago (106-298 MYA) (Johnson et al. 2018) likely after the divergence of the lineage leading to the Aphidomorpha (Adelgids, Phylloxera and Aphids) from the lineage leading to the Coccoidea (scale Insects). We show that both *dnmt3* paralogs are expressed in the ovaries and developing embryos of the viviparous asexual pea aphid. Using an inhibitor of the DNA methylation machinery, 5-azacytidine, we demonstrate that DNA methyltransferases have key roles in oogenesis and embryogenesis in the pea aphid.

## Methods

### Aphid husbandry

Pea aphids (*Acyrthosiphon pisum*), clone N116 (Kanvil et al. 2014) were maintained on broad bean (*Vicia faba*) plants in a Panasonic growth chamber (352H-PE) under long-day conditions, 16L:8D at 20°C.

### Phylogenetic analysis of *dnmt3a* and *dnmt3x*

Pea aphid *dnmt3a* (NCBI, XP_016662566.1) and *dnmt3x* (NCBI, XP_029348651.1) sequences) were used to search for homologs in the genomes and transcriptomes of 48 insect species (Supplementary Table 1), using BLASTp and tBLASTn (using an E-value cut-off of E< 1 × 10^-20^) (Camacho et al. 2009; Altschul et al. 1990). Target databases were NCBI’s protein and nucleotide collection as well as the transcriptome shotgun assembly databases, AphidBase (https://bipaa.genouest.org/is/aphidbase/) and i5k (https://i5k.nal.usda.gov/webapp/blast/, (Poelchau et al. 2015).

Protein sequences were aligned using Clustal Omega (Sievers et al. 2011). Gblocks (Talavera and Castresana 2007) was then used to remove poorly aligned regions, using a maximum number of contiguous non-conserved positions of 20, and otherwise default parameters. The Gblocks output was then realigned using Clustal Omega (Sievers et al. 2011). MrBayes (Huelsenbeck and Ronquist 2001) was then used to carry out Bayesian inference by Markov chain Monte Carlo simulation, using a Poisson amino acid model, a burninfrac value of 0.25, a samplefreq value of 500 and 4 million iterations, mixed models and default priors. Consensus trees were visualised using Figtree (Rambaut 2010).

### Dissection and fixation

Adult asexual pea aphids were collected from plants the day of dissection and ovaries were dissected under a dissecting microscope in cold Phosphate Buffered Saline (PBS) using fine forceps. Tissue was collected into microcentrifuge tubes containing PBS on ice and fixed for 1 h in a mix of 50:40:10 PBS:heptane:formaldehyde (37%) on a nutator. The solution was removed and replaced with ice cold methanol, following two washes. Tissue in methanol was subsequently stored at -20 °C.

### Hybridisation Chain Reaction (HCR)

Immediately preceding hybridisation chain reaction (HCR), tissue was rehydrated over a methanol series: 3:1, 1:1, then 1:3 methanol:PTw (0.3% Tween-20 in PBS) (all PTw used in subsequent steps was 0.3% Tween-20) then rinsed three times in PTw on a nutator at room temperature for 10 minutes each step. Samples were re-fixed in 4 % formaldehyde in PTw for twenty minutes on a nutator and then rinsed in PTw on a nutator for 10 minutes three times. To permeabilise the tissue, samples were then nutated for 45 minutes in detergent solution (1% SDS, 0.5% Tween-20, 50 mM Tris-HCL (pH 7.5), 1 mM EDTA (pH 8.0), and 150 mM NaCl) at room temperature.

HCR was performed according to Molecular Instruments instructions (Choi et al. 2018), with some modifications. Permeabilised tissue was pre-hybridized in 150 µl HCR probe hybridization buffer (Molecular Instruments Inc, 2.4 M Urea, 5 x sodium chloride sodium citrate (SSC), 9 mM citric acid (pH 6.0), 0.1% Tween 20, 50 μg/mL heparin, 1 X Denhardt’s solution, 10% dextran sulphate) at 37 °C for thirty minutes. Probe solution was prepared by adding 0.8 pmol of each sequence specific probe set to 200 µl probe hybridization buffer (Molecular Instruments Inc.) at 37 °C. Pre-hybridization solution was replaced with the prepared probe solution and incubated overnight (16 – 18 h) at 37 °C. Excess probe was removed by washing four times, each for fifteen minutes in 100 µl pre-heated probe wash buffer (Molecular Instruments Inc, (2.4 M Urea, 5X SSC, 9 mM citric acid (pH 6.0), 0.1% Tween, 50 μg/mL heparin)) at 37 °C. Tissue was then washed three times for five minutes in 500 µl 5 x SSCT (0.1 % Tween-20 in 5 x SSC) at room temperature on a nutator. Samples were incubated with 100 µl amplification buffer (Molecular Instruments Inc, 5X SSC, 0.1% Tween 20, 10% dextran sulphate) for thirty minutes at room temperature. Simultaneously, 6 pmol of hairpin h1 and 6 pmol of hairpin h2 (2 µl of a 3 µM stock, Molecular Instruments Inc) were heated to 95 °C for 90 seconds in a thermal cycler, and allowed to cool over thirty minutes protected from light at room temperature. Hairpins (B2-546 for *Ap-dnmt3a*, B3-546 for *Ap-dnmt3x,* B3-546 for *Ap-wg, and* B4-647 for *Ap-vasa*; Molecular Instruments Inc) were added to samples and they were incubated overnight in the dark at room temperature.

The hairpin solution was removed, and excess hairpin was removed by nutating with 500 µl 5 x SSCT at room temperature in the dark, twice for five minutes. Samples were washed twice more with 5 x SSCT as above for thirty minutes each wash, with the first wash including 5 µg/ml DAPI (4′,6-diamidino-2-phenylindole; ThermoFisher Scientific) to stain nuclei. The two 30 min washes were followed by a final five-minute wash. All solution was then replaced with 1 ml 70 % ultrapure glycerol (ThermoFisher Scientific) and samples were stored protected from the light at 4 °C for at least 12 hours prior to imaging. Tissue was mounted on glass slides in a small amount of 70% glycerol with a drop of SlowFade Diamond Antifade Mountant (ThermoFisher Scientific) and stored in the dark until imaging by confocal microscopy. Images were acquired within a week of carrying out HCR, using a Zeiss LSM 880 upright confocal microscope (Zeiss) and the Zen Black software (Zeiss). Images were processed using Zen Blue (v3.1, Zeiss).

### Chemical inhibition of DNA methyltransferases

Asexual pea aphids were reared on an established artificial diet (Kunkel 1977; Prosser and Douglas 1992; supplementary table 2). Artificial diet was stored in single use aliquots at -20°C.

For treatment, preparations of artificial diet were thawed and supplemented with 5-azacytidine to a final concentration of 50 µM (Acros Organics) or an equivalent amount of DMSO for the control diet. 50 µM 5-azacytidine did not cause significantly increased mortality compared to DMSO or water (supplementary Fig. 1). 10 µl of artificial diet was transferred into each of the eight wells of a sterile PCR tube cap strip. These strips were then pushed gently liquid side down into strips of parafilm that had been sterilised with 70 % ethanol, allowed to dry, then stretched over a 70 % ethanol sterilised 96-well PCR tube holder to enable the aphids to feed. Fourth instar (L4 nymph) parthenogenetic pea aphids were collected immediately prior to initiation of feeding. L4 nymphs were identified based primarily on cauda length relative to width at its base (Diamond and Levitis 2016). L4 aphids were transferred individually to the wells of a transparent 96-well microplate. The parafilm-strip cap complexes were then carefully removed and transferred to the 96-well microplates containing aphids, enclosing the aphids. Aphids were maintained in LD conditions (16L:8D, 20°C, 70% RH). The artificial diet solution was replaced daily by preparing fresh parafilm-strip cap complexes. Before replacing the solution, aphids were dislodged from the parafilm by gently stroking with a paintbrush, so they were able to withdraw their stylet. After six days of feeding on supplemented artificial diets, aphids were dissected and processed (as described above) and ovaries were used in HCR (as described above), using *Ap-vasa* (NCBI XM_001948573.5) and *Ap-wingless (Ap-wg;* NCBI XM_001945260.5) probes (Molecular Instruments Inc). Additionally, a second group of six-day fed aphid ovaries were stained with Alexa Fluor phalloidin-488 (Cell Signalling Technologies) and DAPI (ThermoFisher Scientific). Phalloidin staining was carried out by first fixing ovaries in 50:40:10 PBS:heptane:formaldehyde (37%) for forty-five minutes, then washing for five minutes in PTw (0.3 % Tween-20) five times and incubating with PTw (0.3% Tween-20) for one hour. After incubation, ovaries were washed for five minutes with PTw (0.3 % Tween-20) and a solution containing 10 μl Alexa fluor 488 phalloidin (6.6 μM stock, Cell Signalling Technologies) in 200 μl total volume PTw (0.3% Tween-20) was applied. Ovaries were then incubated in the dark for forty-five minutes, after which 800 μl PTw (0.3 % Tween-20) and 1 μl DAPI were added to the ovaries in solution, and ovaries were then incubated for a further thirty minutes in the dark. Following, ovaries were washed as before three times, and then once for five minutes in PBS. Ovaries were stored in 70 % glycerol before mounting in a small amount of 70% glycerol with a drop of SlowFade™ Diamond Antifade Mountant (ThermoFisher Scientific) and imaged within two days using a Zeiss LSM 880 upright confocal microscope.

To understand the effects of this chemical treatment on the life history of this population, a group of aphids were fed azacytidine supplemented or DMSO-control diets for six days. Aphids were then moved to leaf-agar plates (one *Vicia faba* leaf cut close to the petiole using a sterile razor blade embedded in 5 ml of 2% agar supplemented with 1 g/L Miracle-Gro (Miura et al. 2003) and 0.03% Methyl 4-hydroxybenzoate (to inhibit fungal growth) in a 55 x 15 mm petri dish) and monitored for another 14 days (20 days total) and their survival, offspring production, offspring phenotype, and offspring survivability were assessed daily. No more than ten aphids (the focal aphid plus nine offspring) were kept on any one leaf-agar plate at a time, with progeny being spread across several plates where necessary.

### Data analysis

Statistical analyses were performed and graphs were produced using R version 4.1.2. A Cox proportional hazard model was run to assess differences in lifespan between 5-azacytidine and DMSO control focal aphids using the *survival* package. Differences between 5-azacytidine and DMSO control aphids in their daily reproductive output were assessed by fitting a zero-inflated Poisson GLMM fitted with a log link function, and with aphid ID as a random effect and treatment as the fixed effect. This was followed by performing a likelihood ratio test (LRT) comparing the full model to a null model with the fixed effect dropped (using the lmtest package); data was truncated to remove entries prior to initiation of reproduction by a given aphid (all only being able to be 0). Both time from L4 to reproduction and production of disturbed (stillborn) nymphs by 5-azacytidine and DMSO control aphids were assessed by Wilcoxon rank sum test with continuity correction. General survival of nymphs (produced between days six and twenty) were compared by Wilcoxon rank sum test. And fecundity of the offspring of focal aphids (collectively, as pooled groups of offspring) was compared between groups by running a linear model, with aphid ID and day as random effects and treatment as a fixed effect, then performing an LRT as above.

## Results

### *Dnmt3a* and *dnmt3x* arose from an ancient Aphidomorpha specific duplication

Aphid genomes generally show a high number of gene duplications (International Aphid Genomics 2010; Li et al. 2019b; Mathers et al. 2017), but many of these are lineage-specific or species-specific duplications (Mathers et al. 2017; International Aphid Genomics 2010). To investigate the evolutionary history of the duplication event giving rise to *Ap-dnmt3a* and *Ap-dnmt3x* and identify the conserved paralog of the gene, we first identified homologs of *dnmt3* in the genomes and transcriptomes of 48 insect species (Supplementary Table 1).

Dnmt3 protein sequences were aligned, and a phylogeny constructed using Bayesian methods (Ronquist and Huelsenbeck 2003, Fig. 1A) robustly separates these homologs into two clades. One clade contains Ap-Dnmt3a and all other insect Dnmt3 orthologs, while the second clade contains Ap-Dnmt3x and orthologs from eight other aphid species. This separation is consistent with Ap-Dnmt3a being the more highly conserved, possibly ancestral, copy of Dnmt3, while Ap-Dnmt3x is more diverged.

**Figure 1.**
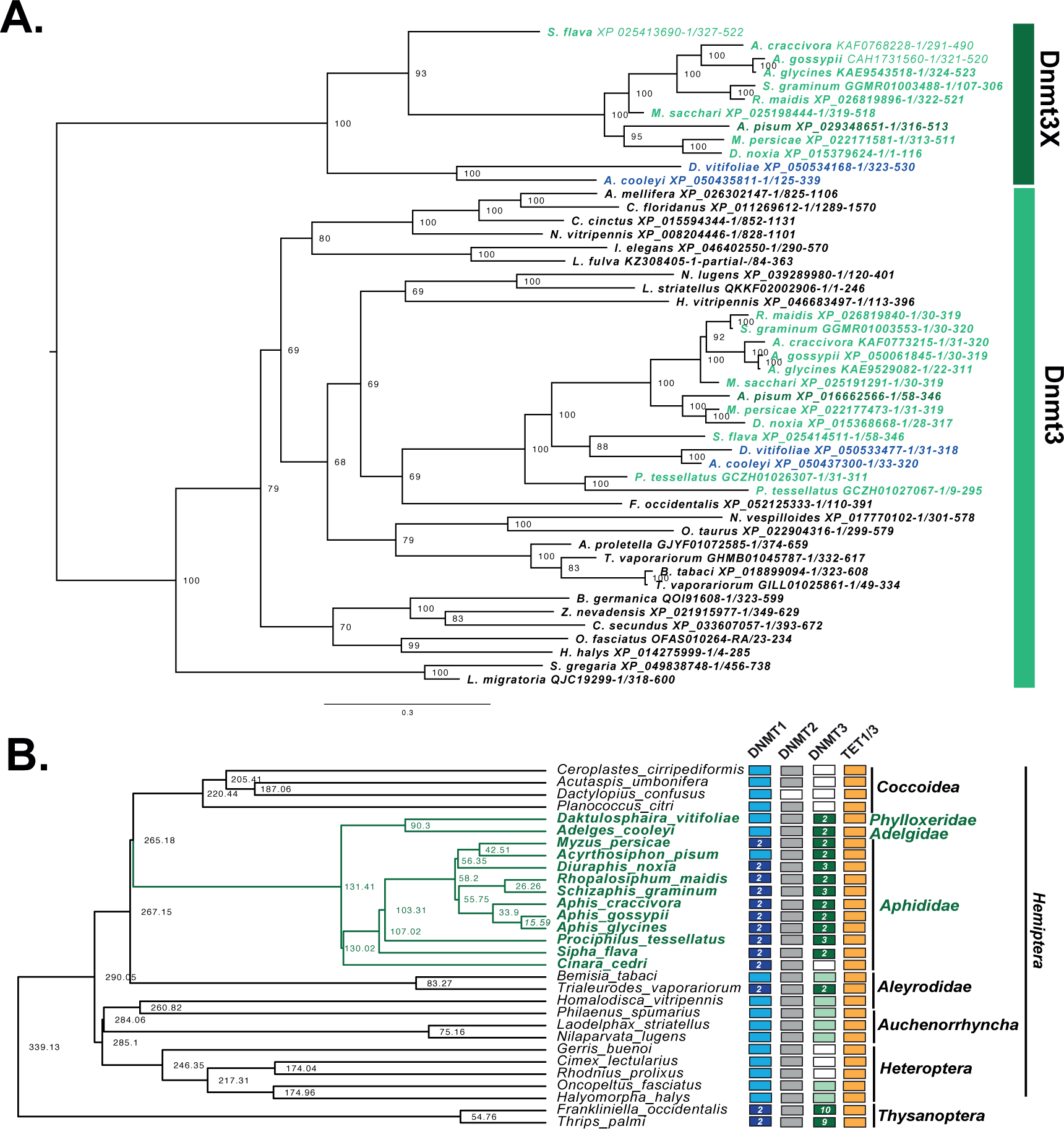
Evolutionary History of the Dnmt3 paralogs. **A)** Bayesian phylogeny of insect Dnmt3 proteins from 48 insect species. The phylogenetic tree is mid-point rooted and posterior probabilities are shown at nodes. This analysis reveals two well supported clades, an ancestral clade corresponding to the conserved Dnmt3 protein (light green box) containing orthologs from a wide phylogenetic range of insect species, and the second containing the dnmt3 paralogs (dark green box). Aphid species are shown in green text while Adelogidea and Phylloxeroidea are shown in blue text. Full species names and protein accessions are provided in Supplementary File 1. **B)** Conservation of genes involved in DNA methylation across Hemiptera and Thysanoptera. The phylogenetic relationships between species and divergence times (shown at nodes) are based on TimeTree of Life (Kumar et al. 2022). Lighter colours indicate the presence of a single homolog in the genome, while darker colours indicate the presence of paralogs (the number of paralogs is indicated). *dnmt1*, *dnmt2*, and *tet1* are generally very well conserved across the Hemiptera and Thysanoptera although duplicates of *dnmt1* are often seen, which may indicate the potential for neo or subfunctionalisation of these paralogs. In contrast, *dnmt2* and *tet1* are never duplicated. *dnmt3* exhibits more variability (full figure showing a wider range of insect species is shown in Supplementary Fig. S2).

The Dnmt3x clade also includes sequences from the grape phylloxera (*Daktulosphaira vitifoliae*), a representative species of the Phylloxera, and the gall adelgid (*Adelges cooleyi*), a representative of the Adelgidae. The Phylloxera and Adelgidae diverged from the Aphididae approximately 106 million years ago (Johnson et al. 2018). The conservation of both paralogs in these species indicates that the duplication giving rise to *dnmt3a* and *dnmt3x* is ancient and likely occurred between 106 and 298 million years ago (Johnson et al. 2018). This gene duplication likely occurred after the divergence of the lineages giving rise to the Coccoidea, where *dnmt3* has been lost (Fig. 1B), and the Aphidomorpha (Adelgoidea, Phylloxeroidea and Aphidoidea), all of which have at least two copies of *dnmt3*: *dnmt3a* and *dnmt3x*. However, it is also possible that the duplication occurred prior to this in the lineage leading to the Coccoidea/Aphidomorpha after the divergence of the lineage leading to the Aleyrodidea (whiteflies) 338 million years ago (Johnson et al. 2018), and both duplicates were subsequently lost in the lineage leading to the Coccoidea. Either way these paralogs have been stably maintained in the genomes of these species for at least 106 million years. Although the two *dnmt3* paralogs are present on the same chromosome in pea aphid (X) they are separated by 151 protein coding genes and 6.19 Mbp, and there was no detectable synteny even between closely related species (Supplementary Fig. 3), consistent with this duplication being ancient.

The dnmt3a paralog identified in Aphidomorpha is clustered with the other insect dnmt3 orthologs (Fig. 1) suggesting it may be more similar to the ancestral gene copy (Fig. 1). Analysis of domain architecture (Supplementary Fig. S4) and intron exon structure (Supplementary Fig. S5) shows that proteins in the Dnmt3x paralog group retain more organisational similarity to the Orthopteran Dnmt3 protein sequences, which may suggest subfunctionalisation of the ancestral gene between these two paralogs (Supplementary Fig. S4, S5).

### *Ap-dnmt3a*, *Ap-dnmt3x*, and *Ap-vasa* mRNAs colocalize in the germ cells of developing embryos and the germaria

To examine possible subfunctionalization of Dnmt3s, we investigated the expression of *Ap-dnmt3a* and *Ap-dnmt3x* during development with mRNA localisation using *in situ* hybridization chain reaction (HCR). Expression of both *dnmt3* paralogs consistently colocalises with *Ap-vasa* (a conserved marker of germ cells (Chang et al. 2007; Chang et al. 2006) indicating that these *dnmt3* paralogs are expressed in the developing germ cells of the pea aphid (Fig. 2,3).

**Figure 2.**
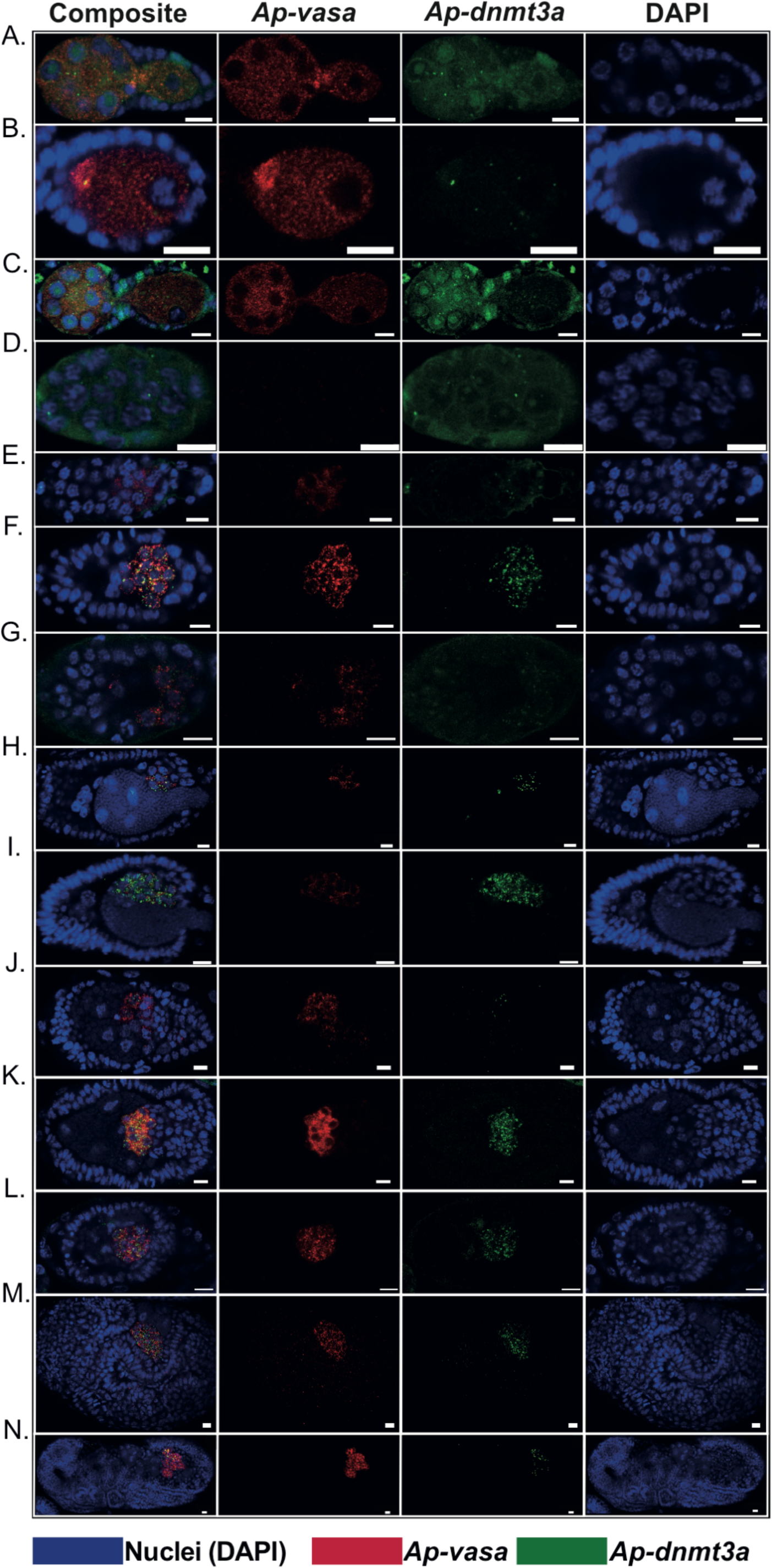
Expression of *Ap-vasa and Ap-dnmt3a* during viviparous oogenesis (A-C) and embryogenesis (D-N). <u>A-C oogenesis</u>: *Ap-vasa* and *Ap-dnmt3a* are maternally provided and mRNAs are detected in the germarium (A) and at stage 0 (B) through to stage 1 of oogenesis (C). *Ap-vasa* mRNA is strongly localised to the anterior of the developing oocyte (as has been previously reported (Chang et al. 2007), however, this is not observed for *Ap-dnmt3*. <u>D-N</u> <u>embryogenesis:</u> Between late oogenesis and early embryogenesis (Stage 3, D) maternal mRNAs are cleared from the embryo and no signal for either *Ap-vasa* nor *Ap-dnmt3a* can be detected. Embryonic expression of both *Ap-vasa* and *Ap-dnmt3a* is detected early in stage 6 after the cellular blastoderm has formed (E), expression of both genes intensifies during mid (F) to late stage 6 / early stage 7 (G) and is localised to the presumptive germ cells. The co-localisation of *Ap-dnmt3* with *Ap-vasa* persists throughout the rest of development, including as the germ cells migrate dorsally to accommodate the invading endosymbiotic bacteria at stage 7 (H,I). At the beginning of gastrulation (Stage 8, J), the bacteria have finished invading and the germ band begins to invaginate (arrowhead). Stage 9 germ band invagination (K-L), stage 11 S-shaped embryo (M) and post-katatrepsis as the germ band retracts at stage 17 (N). Scale bars represent 10 μm.

**Figure 3.**
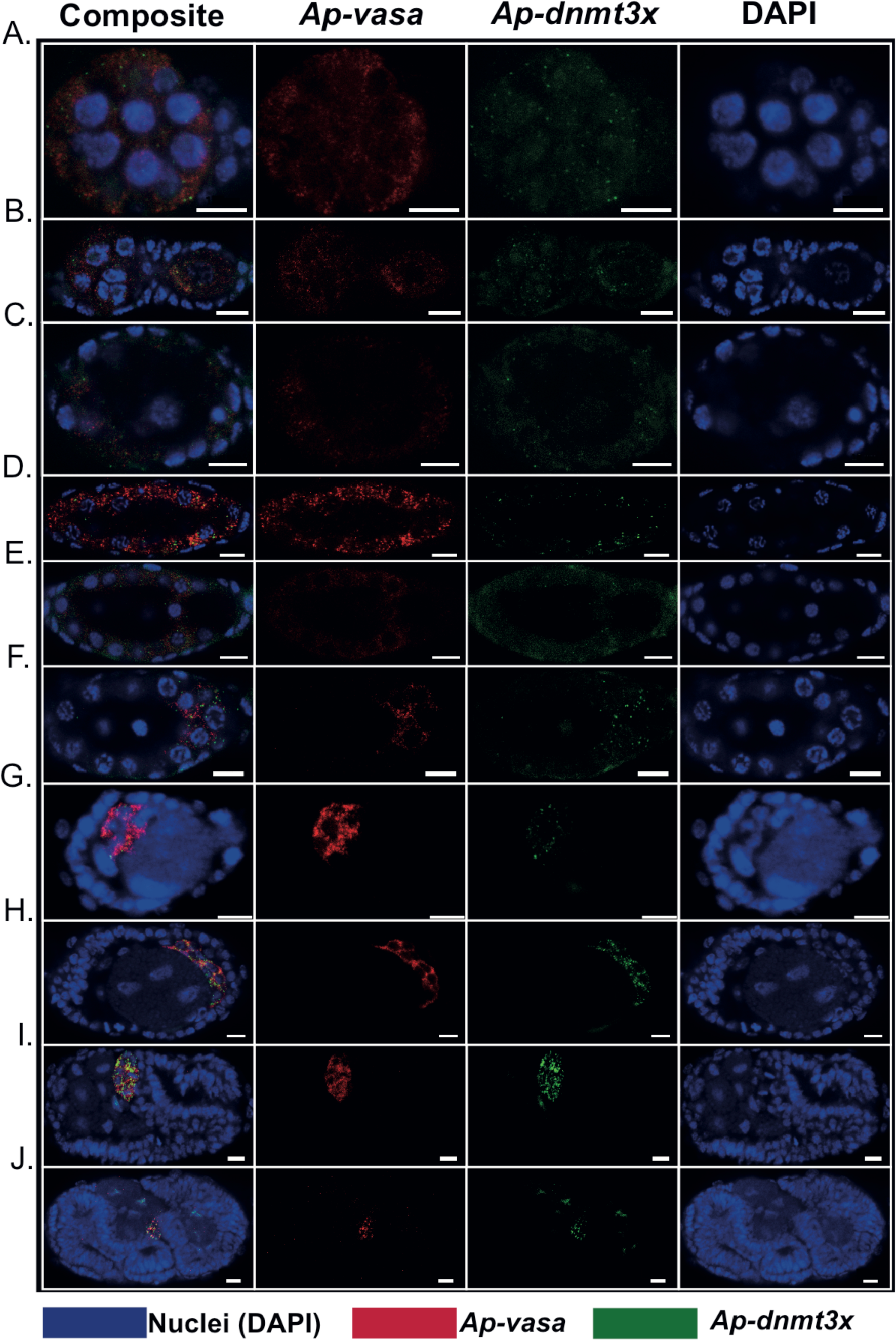
Expression of *Ap-vasa and Ap-dnmt3x* during viviparous oogenesis (A-B) and embryogenesis (C-J). <u>A-B oogenesis</u>: *Ap-vasa* and *Ap-dnmt3a* are maternally provided and mRNAs are detected in the germarium (A) and at stage 1 of oogenesis (B), where *Ap-vasa* mRNA and *Ap-dnmt3x* mRNAs are strongly localised to the anterior of the developing oocyte (as has been previously reported for *Ap-vasa* (Chang et al. 2007). <u>C-J embryogenesis:</u> Between late oogenesis and early embryogenesis (Stage 2, C) maternal mRNAs are cleared from the embryo and no signal for either *Ap-vasa* nor *Ap-dnmt3x* can be detected. Embryonic expression of *Ap-vasa* is detected ubiquitously throughout the cellular blastoderm has formed (stage 5, D), while punctate expression of *Ap-dnmt3x* is detected in cells of the posterior blastoderm only (D). As the germ cells are specified (stage 6 (E,F)), expression of both genes intensifies and is localised to the presumptive germ cells. The co-localisation of *Ap-dnmt3x* with *Ap-vasa* persists throughout the rest of development, including as the germ cells migrate dorsally to accommodate the invading endosymbiotic bacteria at stage 7 (G). At the beginning of gastrulation (Stage 8, H), the bacteria have finished invading and the germ band begins to invaginate (arrowhead). Stage 10 Germ band invagination (I) Stage 11 S-shaped embryo (J). Scale bars represent 10 μm.

*Ap-dnmt3a* (Fig. 2A,B) and *Ap-dnmt3x* (Fig. 3A,B) mRNAs are both maternally provided and are weakly detected in the germaria and in early oocytes (stages 0, 1). *Ap-vasa* localises to the anterior of the developing oocyte at stage 1 (oocytes have just been pinched off from the germarium by the follicle cells, but are still connected to the germarium by the trophic core) (Fig. 3B) and we see similar localisation for *Ap-dnmt3x* (Fig. 3B), but only weakly for *Ap-dnmt3A* (Fig. 2B). *Ap-vasa*, *Ap-dnmt3a* and *Ap-dnmt3x* mRNAs are not detected at stages 2-3 of development (Fig. 2C,D; Fig. 3C), likely as a result of clearance of maternal mRNAs. *Ap-vasa* mRNA is not detected again until stage 5, where it is located in the cytoplasm surrounding a subset of nuclei once they have localized to the periphery, presumed to be the presumptive germ cells (Fig. 3D). At the same stage, before the germ cells are specified at the posterior end of the embryo, punctate expression of *Ap-dnmt3x* mRNA is detected around the nuclei around the embryo’s periphery, across the embryo but with greater signal intensity at the posterior end where the presumptive germ cells are located (Fig. 3D). At stage 6, the germ cells are specified and split the embryo into a central syncytium and posterior syncytium. From this stage onwards *Ap-vasa* mRNA is restricted to the germ cells, and *Ap-dnmt3a* and *Ap-dnmt3x* mRNAs co-localize with high-specificity with *Ap-vasa* as the germ cells migrate as the result of infiltration of the endosymbiotic bacteria at stage 7 (Fig. 2H,I; Fig. 3G) and as the germ band invaginates at stage 8/9 (Fig. 2J-L, Fig. 3H). This co-expression persists throughout mid-late development (Fig. 2M,N; Fig. 3I,J). The tight temporal and spatial colocalistion of *Ap-dnmt3a*, *Ap-dnmt3x* with *Ap-vasa*, which has a known and conserved role in germ-cell specification, may indicate that the *Ap-dnmt3* genes may also have a role in this process.

### Treatment with the methylation inhibitor 5-azacytidine treatment causes defects in reproduction and development in viviparous pea aphids

To examine potential roles of DNA methylation in pea aphid reproduction and development we treated aphids with the methylation inhibitor 5-azacytidine. We found that while having equal lifespans (Cox proportional hazards model, p = 0.7, Supplementary Fig. S1), and exhibiting no difference in the first day of reproduction (5-azacytidine mean = 3.71 days, DMSO mean = 4.2 days, Wilcoxon rank sum test: W = 25.5, p = 0.3531, Fig. 4A), 5-azacytidine treated aphids produced significantly fewer nymphs per day than DMSO-control aphids over a twenty day period (GLMM, LRT: χ^2^ = 8.7787, p = 0.003048, df = 1; treatment: z = 3.828, p = 0.000129; Fig. 4B).

**Figure 4.**
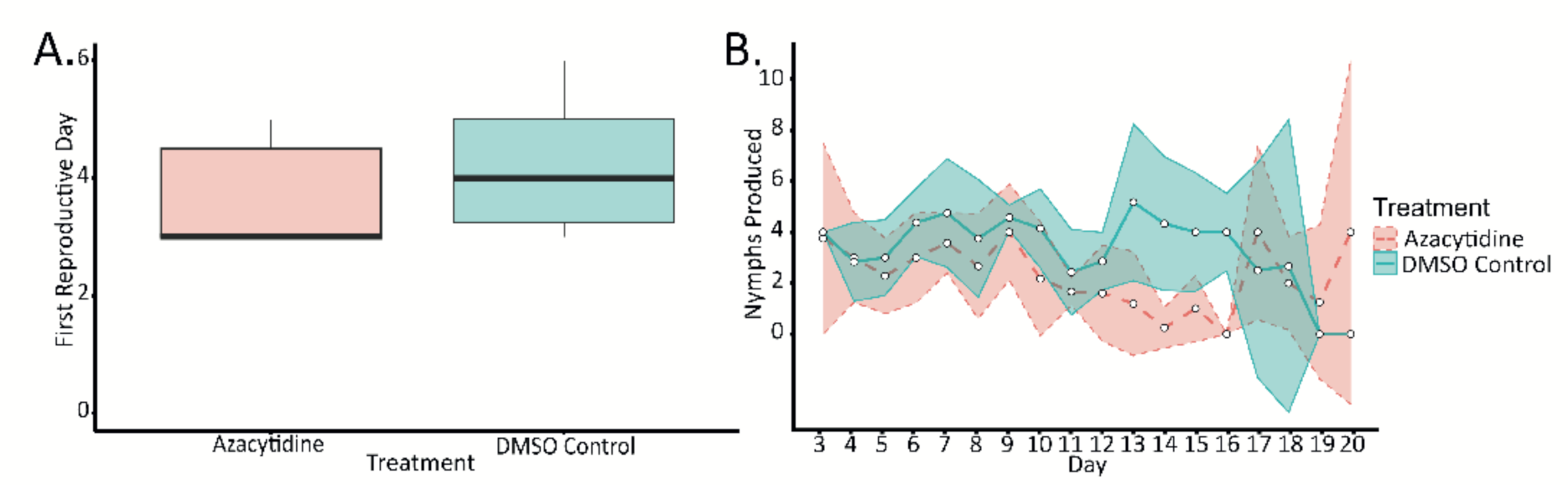
5-azacytidine treatment alters reproductive rate in viviparous pea aphids. **A.** Boxplots showing the first day reproduction was observed for 5-azacytidine and DMSO treated groups. Reproduction initiated, at the earliest, 3 days after the onset of treatment. **B.** Daily reproductive output of 5-azacytidine and DMSO treated aphids, from day 3 (the earliest day nymphs were recorded for any aphid), until day 20 (at which point the experiment was concluded due to sample size). Points represent mean nymphs produced by all individuals of a treatment and ribbons indicate 95 % confidence intervals.

Treatment with 5-azacytidine resulted in the production of ‘stillborn’ nymphs, a phenotype that was never seen in the DMSO controls. Stillborn nymphs were either dead at birth or died shortly after birth. The gross morphology of stillborn nymphs was normal, and the only obvious defect seen was an inability to extend their legs, perhaps owing to a failure of the serosa to rupture (Fig. 5A). 75% of all azacytidine treated aphids produced at least one stillborn nymph (Fig. 5A) and no stillborn nymphs were observed in the DMSO control (azacytidine treated aphids produced significantly more stillborn nymphs than did DMSO-control aphids, Wilcoxon rank sum test, W = 3995, p = 0.001441; Fig. 5A). Outside of the production of stillborn nymphs, all nymphs produced by 5-azacytidine treated adults and DMSO-control adults appeared morphologically normal, and there was no difference in the mortality of the offspring of azacytidine treated and control aphid nymphs (proportion of total nymphs dead, Wilcoxon, W = 49, p = 0.1874), or the reproductive output of these offspring (LM, LRT: χ2 = 0.8084, p = 0.3686, df = 3, 2; Supplementary Fig. 6). Stillborn nymphs were also not observed among the offspring of the offspring of treated aphids.

**Figure 5.**
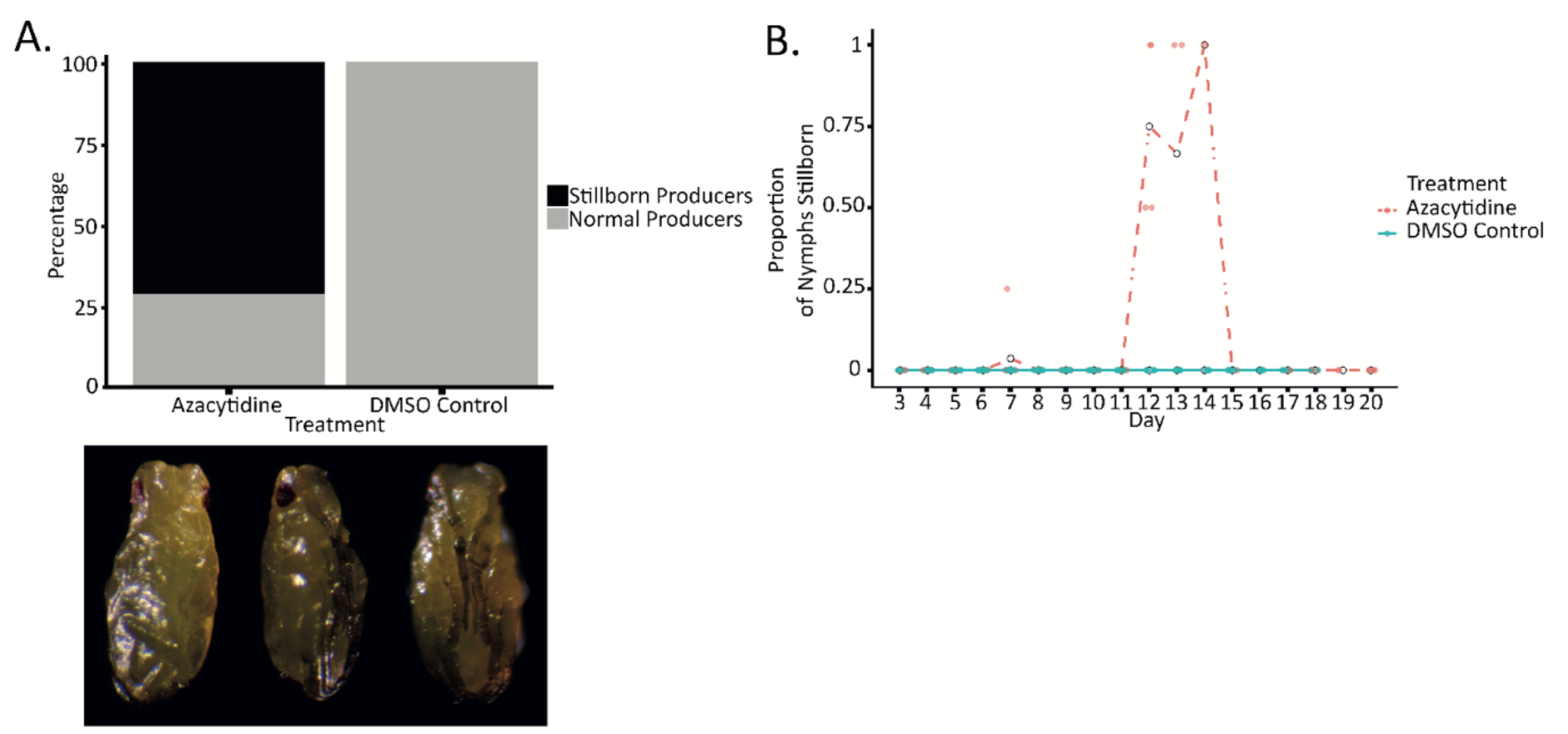
5-azacytidine treatment causes mature embryos to be stillborn in viviparous pea aphids. **A.** Percentage of all 5-azacytidine (50 µM) treated and control (DMSO) aphids that produced ‘stillborn’ (bottom panel: stillborn nymphs were either dead at birth or shortly after birth and their gross morphology was normal) aphids after treatment from day 0 to day 6. Stillborn offspring were only produced by 5-azacytidine treated aphids. **B.** Daily mean proportion of stillborn nymphs for each 5-azacytidine treated and control aphid, between day 3 (the first day nymphs were observed) and day 20.

Each aphid that produced stillborn nymphs had between one and four total stillborn nymphs between days twelve and fourteen, but mostly appearing on day twelve (six days after the cessation of treatment) (Fig. 5B). Temporally this coincided with the period of reduced reproduction seen in 5-azacytidine treated aphids (Fig. 4B). It is interesting to note that, while nymph production slowed down during the production of stillborn nymphs, it generally did not stop completely. And, where these stillborn nymphs appeared more than once during this period, they did so in chains uninterrupted by normal nymphs (except for one stillborn nymph produced by a single aphid on day 7), suggesting that 5-azacytidine treatment was affecting a critical period in development essential for embryo maturation 6-8 days prior to birth, or that 5-azacytidine has a high level of stability in the pea aphid.

The decreased reproductive rate in 5-azacytidine treated aphids (Fig. 4B) may indicate defects in early development that result in resorption of the oocyte or embryo (Ward and Dixon 1982). To examine this, we dissected and stained ovaries with developing embryos with phalloidin and DAPI to identify gross morphological differences (Fig. 6) and performed HCR with *Ap-vasa* and *Ap-wg* to identify specific defects in germ-cell specification and segmentation respectively (Fig. 7).

**Figure 6.**
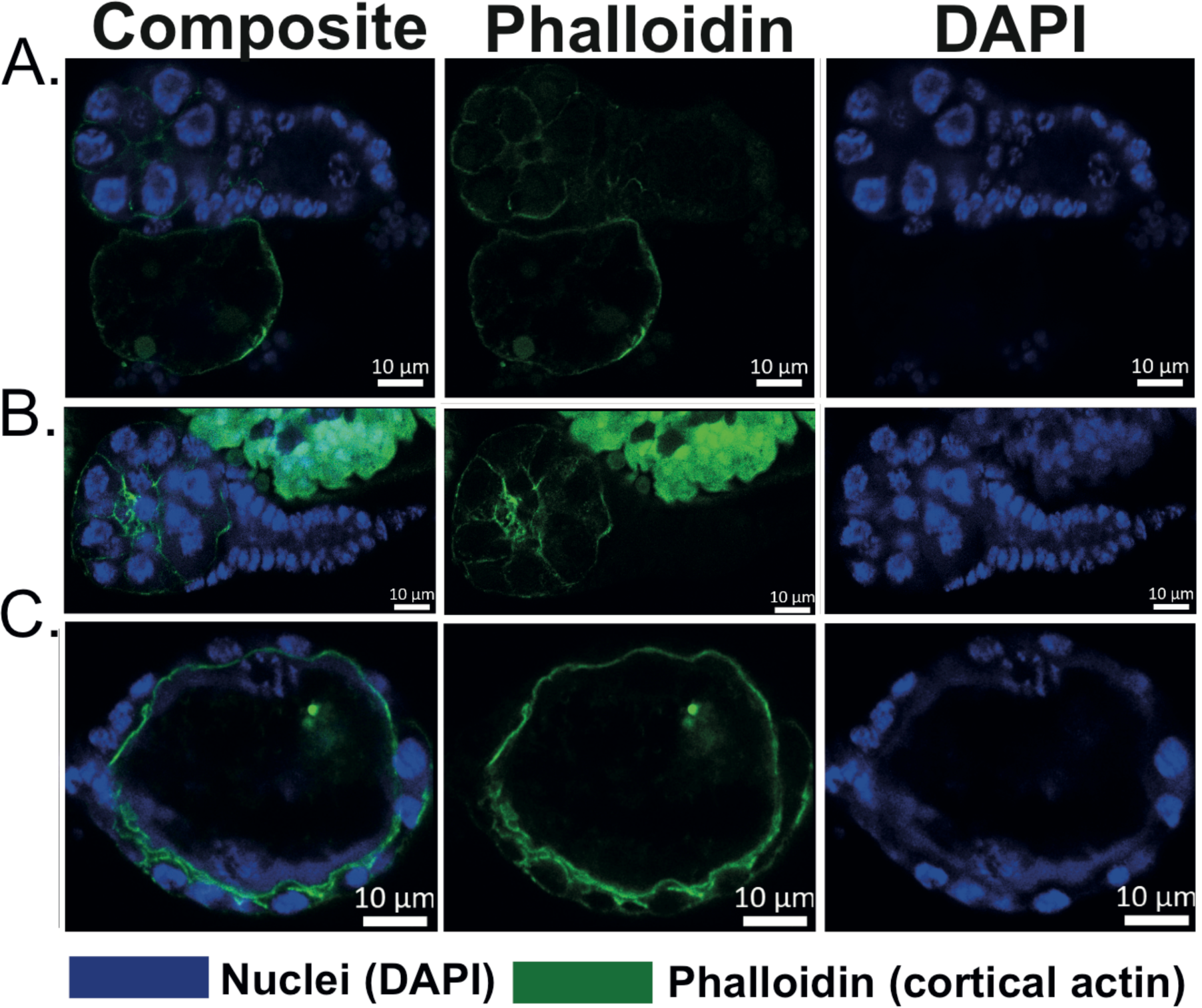
5-azacytidine treatment causes early defects in oogenesis in viviparous pea aphids. Representative images of *A. pisum* ovary samples stained with phalloidin (stains actin, green, middle) and DAPI (stains nuclei, blue, right) after aphids were treated with 5-azacytidine, a disruptor of DNA methyltransferases. Images are single plane of focus per row, and individual channels are presented alongside composite (left). **A.** Phalloidin staining reveals an unusual structure, positive for actin and appearing ectopic to the germarium and oocyte, possibly a mass of cortical actin. **B.** The second row shows two adjacent oocytes and the germarium, the oocytes appear odd in that they do not appear to have separated from each other, and form a long continuous chain. **C**. The third row shows an oocyte with an odd size and morphology for the number of nuclei. Scale bars represent 10 μm.

**Figure 7.**
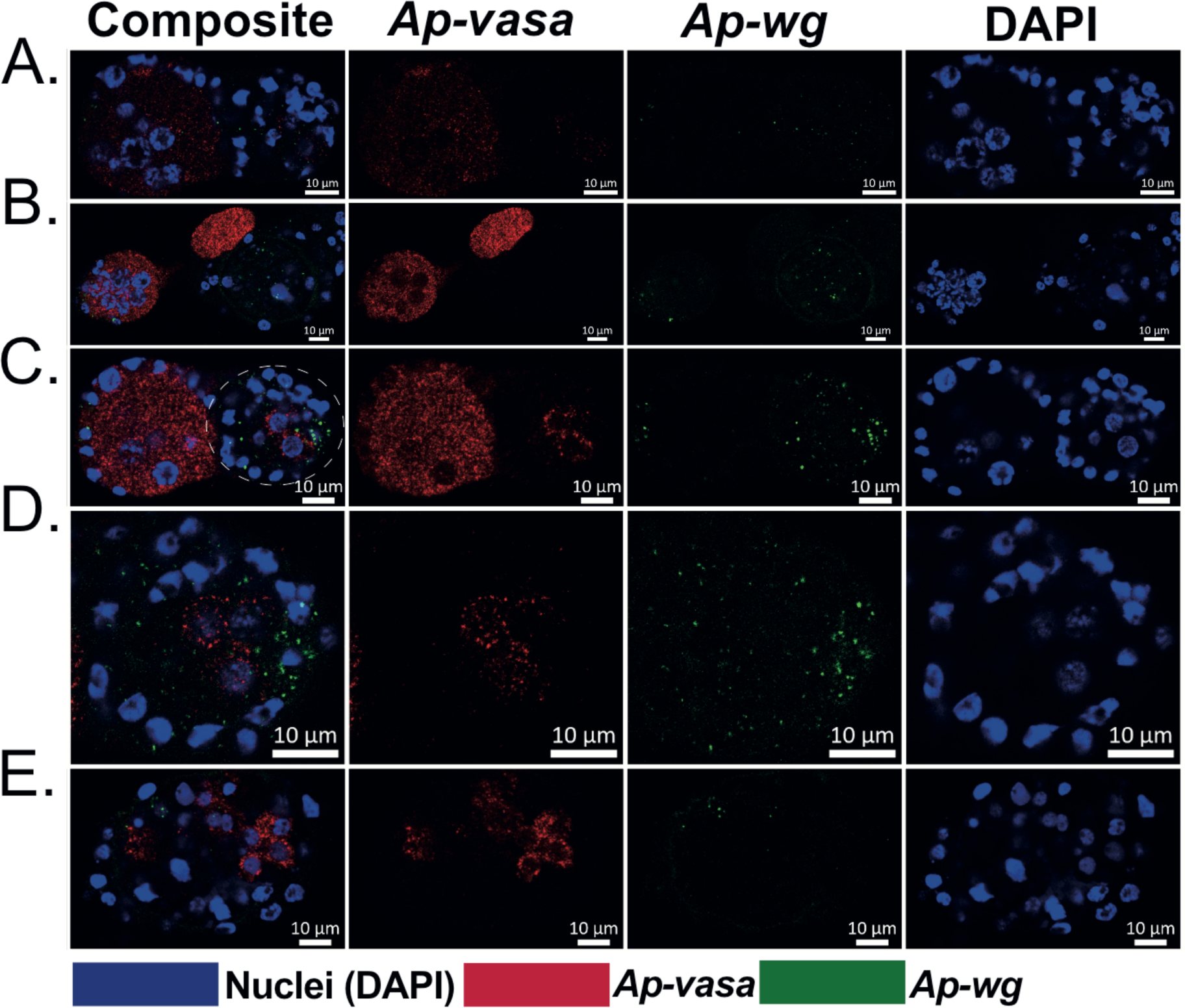
5-azacytidine treatment causes defects in early embryogenesis in viviparous pea aphids. Representative images of *A. pisum* ovary and embryo samples showing expression of *Ap-vasa* (red), *Ap-wg* (green) DAPI (stains nuclei, blue, right) after aphids were treated with 5-azacytidine, a disruptor of DNA methyltransferases. Images are single plane of focus per row, and individual channels are presented alongside composite (left). Due to the severity of defects embryos are difficult to accurately stage. A) a malformed germarium and early oocyte (right) and a misshapen early blastoderm embryo (left), both specimens show aberrant expression of *Ap-vasa* which is notably absent from the blastoderm stage embryo and suggests that germ cells have not been correctly specified. B). Note the unusual arrangement of cells in the germarium (left), the embryo to the right is consistent with an early blastoderm stage embryo (stage 5) but has fewer cells. Note also the extrusion of cytoplasm from the embryo that is positive for *Ap-vasa* mRNA. C). Shows an oocyte/early embryo with two nuclei and a polar body visible (left), we see no localisation of *Ap-vasa* mRNA, to the right the embryo has too few cells and is round rather than elongated, yet we see expression of *Ap-vasa* and *Ap-wg* consistent with a late blastoderm (stage 6 embryo), enlarged in D. E). Here we see defects in *Ap-vasa* expression that we attribute to defects in establishing the anterior-posterior and dorso-ventral axes. Rather than being restricted to the posterior we see *Ap-vasa* expression in cells at the anterior and dorsal surface of the embryo, consistent with these cells being aberrantly specified as germ cells. There is no *Ap-wg* expression at the posterior.. Scale bars represent 10 μm.

After treatment for six days, the germaria, oocytes and very early embryos of azacytidine treated aphids appear unusual. In the germarium, the general morphology was disturbed with defects in the gross structure (e.g. Fig. 6A, where an extrusion of cortical actin is seen originating from the germarium) as well as higher numbers of cells and disorganisation of the cells contained within the germarium (Fig. 6A, B), possibly due to an arrest in the progression of oogenesis. There is a failure of oocytes to separate from the germarium and from each other (Fig. 6B) which may indicate defects in follicle cell differentiation. In other species, such as *Tribolium* (Baumer et al. 2012), differentiation of the follicle cell population is required to establish proper morphology of the egg chamber and separation of the egg chambers, although it has been suggested that aphid follicle cells don’t diversify into distinct populations (Michalik et al. 2013). In severe cases, we see multiple oocytes fail to separate and form a long, continuous chain surrounded by follicle cells around their edges, this phenotype is similar to that seen with *dnmt1* RNAi in *Oncopeltus faciatus* (Amukamara et al. 2020; Bewick et al. 2019; Washington et al. 2021).

Defects in gross morphology (Fig. 6C) and gene expression (Fig. 7) are observed in the early stages of embryogenesis as a result of 5-azacytidine treatment. These defects make the embryos difficult to confidently stage, however, using the expression of *Ap-vasa* as a marker of germ cell specification (Chang et al. 2007) and *Ap-wg* as a marker of posterior specification and segmentation (Duncan et al. 2013a), it is clear that there is a range of phenotypes observed (Fig. 7). We observe defects of cell number and organisation in the germaria (Fig. 7A,B), that corresponds with low levels or abnormal *Ap-vasa* expression. In some cases we see extrusions from the early oocyte/embryo that stain positive for *Ap-vasa* (Fig. 7B). We also see significant defects or delays in early embryogenesis (Fig. 7C, D), with relatively normal arrangement of *Ap-vasa* and *Ap-wg* expression (Fig. 7C, D) but this expression is likely occurring prematurely with less cells than we would expect staining positive for each marker (only two cells to the posterior of the embryo staining positive for *Ap-wg* adjacent to only three cells staining positive for *Ap-vasa*). Additionally, based on the expression of *Ap-vasa* and *Ap-wg* this embryo would be considered late stage 6 (blastoderm), yet the overall morphology and number of cells present in this embryo is inconsistent with a blastoderm stage embryo. More severe defects in embryogenesis as a result of 5-azacytidine treatment are also observed with some embryos displaying completely aberrant expression of *Ap-vasa* and *Ap-wg* (Fig. 7E, with *Ap-wg* being expressed in a small central domain bounded by two *Ap-vasa* expression domains), consistent with defects in axis patterning.

From stage seven onwards no defects in morphology or expression of *Ap-vasa* or *Ap-wg* were observed (data not shown). This may suggest that the activity of the DNA methylation machinery is integral only at the early stages of development, or that the effects later in development are subtle as these embryos have developed beyond a critical period.

## Discussion

Cyclical parthenogenesis is a novel and remarkably successful life-history strategy that is observed in most extant aphids (Dixon 1985; Brault et al. 2010; Guerrieri and Digilio 2008). Cyclical parthenogenesis evolved in the Aphidomorpha approximately 106-298 million years ago (Grimaldi and Engel 2005) from an ancestor that was oviparous (Gavrilov-Zimin 2021) and sexually reproducing (Dixon 2012). This reproductive strategy allows aphid species to switch between viviparous parthenogenesis and oviparous sexual reproduction dependent on environmental cues (Ogawa and Miura 2014). However, the mechanisms that control the plasticity of reproductive mode and ovary development in the offspring are largely unknown. Here, we investigate the evolutionary history and expression of two paralogs of the *de novo* methyltransferase *dnmt3* in the pea aphid. We further characterise the function of the DNA methyltransferase proteins in oogenesis and embryogenesis using a chemical inhibitor, 5-azacytidine.

Duplicated genes have been long thought to be a source of evolutionary novelty (Ohno 1970). Aphidomorpha, including aphids (International Aphid Genomics 2010; Li et al. 2019b; Mathers et al. 2017; Fernández et al. 2020), phylloxera and adelgids (Julca et al. 2020; Li et al. 2023) all have very high levels of gene duplications. Many of these gene duplications are species-specific and therefore likely to be more recent (Fernández et al. 2020), but a burst of duplication in the lineage leading to the Aphidomorpha also means that a significant proportion are ancient (occurring between 106 and 298 million years ago) and conserved, (Julca et al. 2020; Li et al. 2023), and these duplications may have had a role in facilitating the evolution of novel and diverse life history strategies, such as parthenogenesis, within this clade.

Here we show that the duplication that gave rise to two copies of *dnmt3* in the pea aphid (Walsh et al. 2010) likely occurred between 106 and 298 million years ago in the Aphidomorpha (the lineage that gave rise to the Aphididae, Adelgidae and Phylloxeridae). Duplicated genes that are redundant tend to be silenced or removed from the genome (Lynch and Conery 2000), but some duplicates are retained either because they have evolved new functions (neo-functionalization; (Ohno 1970)) or taken on part of the function of the ancestral gene (sub-functionalisation; (Force et al. 1999). Orthologs of *dnmt3*, a gene that is implicated in *de novo* DNA methylation, are found across plants, animals and fungi (Zemach et al. 2010) with the gene evolving ∼1,000 million years ago (Golding 1996). Although it is ancient, the *dnmt3* gene is also evolutionarily labile and has been lost repeatedly in multiple insect lineages, even in closely related species (Duncan et al. 2022; Bewick et al. 2017; Provataris et al. 2018). Therefore, the broad conservation of both *dnmt3* paralogs in Aphidomorpha is consistent with each having an essential function. To begin to address the possible functions of these paralogs we examined the expression of both paralogs in viviparous pea aphid ovaries and during embryonic development (Fig. 2, 3).

We found that *Ap-dnmt3a* and *Ap-dnmt3x* mRNA colocalise temporally and spatially with *Ap-vasa* mRNA, a germline marker (Chang et al. 2006), in asexual pea aphid ovaries and throughout embryonic development. The expression of both *dnmt3* paralogs maternally and in the presumptive germ cells at the same time as *Ap-vasa* may indicate a role for these genes in germ cell specification or maturation. Dnmt3 is a positive regulator of translation, in honeybees (Kucharski et al. 2023b) and mammals (Rona et al. 2016). Dnmt3 has been shown to associate with specific histone modifications through the PWWP domain (Rona et al. 2016; Du et al. 2015; Dhayalan et al. 2010; Kucharski et al. 2023b), which is also present in Ap-Dnmt3x and so it is possible that, in this context, the dnmt3 paralogs may act as regulators of germ cell specification or maintenance through transcriptional regulation.

To explore possible roles for DNA methyltransferases in the ovary, we treated asexual pea aphids with 5-azacytidine which inhibits methyltransferase activity by causing them to irreversibly bind to DNA, where after they are degraded (Stresemann and Lyko 2008). This treatment likely reduces the ability of any DNA methyltransferases to carry out any methylation-independent functions they might have. Although this treatment does not allow us to differentiate between the functions of Dnmt3a, Dnmt3x, or Dnmt1, it can tell us the role, if any, of DNA methyltransferases in reproduction and development of the pea aphid. Exposure to 5-azacytidine led to gross morphological defects in early embryos and oocytes, and disorganisation of germaria, likely because of failures in oocyte specification, separation and production. This phenotype, is broadly consistent with the inhibition/knockdown of DNA methyltransferases in other hemipterans (Dnmt1 in *O. fasciatus* (Washington et al. 2021; Bewick et al. 2019; Cunningham et al. 2024) and *Bemisia tabaci* (Shelby et al. 2023)) and *dnmt1* is maternally expressed in the pea aphid (data not shown). However, some phenotypes observed are different that lead us to suggest they are *dnmt3*-specific. We observe defects in the tropharium and in segregation of the oocyte after 5-azacytidine treatment that are not seen with other hemipteran species after *dnmt1* knockdown. Intriguingly, down-regulation of *dnmt1* in *B. tabaci* has been shown to cause a decrease in the expression of *cdc20* (Cunningham et al. 2024; Shelby et al. 2023), a gene involved in mitosis, it remains possible that *dnmt1* inhibition has an aphid-specific phenotype, but given its very broad and consistent effect in other species we think that unlikely. Differences in the function of DNA methyltransferases between the pea aphid and other hemipteran species could also reflect peculiarities of aphid oogenesis. Parthenogenetic aphids are able to activate their oocytes independently of fertilisation; haploid oocytes are arrested at prophase, and a modified meiosis II division gives rise to a diploid oocyte, aster self-organisation (Riparbelli et al. 2005) facilitates a single maturation division that gives rise to the diploid polar body (Gautam et al. 1993; Le Trionnaire et al. 2008; Blackman 1978). Therefore, it is possible that the phenotypes we observe in the germarium, oocyte and follicle cell populations after 5-azacytidine treatment (Fig. 6,7) are due to aphid specific roles for the DNA methyltransferases in parthenogenetic oogenesis.

We also see defects in embryogenesis and patterning (Fig. 7) as a result of 5-azacytidine treatment, with both premature and aberrant expression of *Ap-vasa* and *Ap-wg* (a marker of posterior cell fate and segmentation) as well as abnormal embryo morphology. These phenotypes may reflect defects in embryonic cell division similar to the defects that we see in oogenesis. Alternatively, in *N. vitripennis* knockdown of *dnmt1* resulted in failure of cellularisation and arrest of embryogenesis at or around gastrulation that is consistent with dysregulation of the maternal-to-zygotic transition (Zwier et al. 2012; Arsala et al. 2022). It is therefore possible that the embryonic defects that we observe are due to similar dysregulation of the maternal-to-zygotic transition. We do also observe patterning defects that may be due to mis-localisation or mis-expression of maternal factors that are required to establish polarity of the embryo and the segmentation cascade (Lynch 2019; Sachs et al. 2015; Pechmann et al. 2021). Consistent with 5-azacytidine treatment causing disruption of the embryonic axes we do observe mis-expression of *Ap-vasa* at the anterior and dorsal surface of some embryos. Axis specification in viviparous aphids may involve novel patterning molecules or mechanisms (Duncan et al. 2013b), and it would be interesting to determine if 5-azacytidine treatment causes similar defects in oviparous reproduction, or if the role of DNA methyltransferases in axis specification is unique to viviparous reproduction in aphids. It has been suggested that germ cell specification relies on germ plasm preformed within the egg prior to cellularisation (Chang et al. 2006), it is possible that aberrant *Ap-vasa* expression is due to defects in this process, but this does not explain the mis-expression of *Ap-wg.* Importantly, we did not observe any defects in patterning or morphology in mid-embryogenesis, this may indicate that DNA methytransferases are only required for critical windows in early viviparous development or may be indicative of resorption of severely affected oocytes and embryos (Ward and Dixon 1982). In a related aphid species early embryos are resorbed, mid-stage embryos are paused and late stage embryos continue development in response to starvation stress, which may indicate the existence of developmental checkpoints at these stages. Our data does indicate a lower reproductive rate between days 13-16 which may be consistent with embryo resorption (Fig. 4).

Treatment with 5-azacytidine also resulted in the production of stillborn nymphs (Fig. 5). This phenomenon was constrained to a relatively small temporal period during treatment, likely corresponding to a small window of development where embryos are sensitive. The appearance of stillborn nymphs over just three days is consistent with aphids younger than the critical period being resorbed or arrested (Ward and Dixon 1982). What defects lead to the stillborn phenotype is unclear, but it could reflect a role for the DNA methyltransferases in processes critical for embryo maturation, such as rupture and withdrawal of the extraembryonic membranes (Horn et al. 2015; Schmidt-Ott and Kwan 2016). However, the serosa of the parthenogenetic pea aphid is reduced compared to other hemipteran species (Miura et al. 2003) and the role of the extraembryonic membranes in viviparous aphid development isn’t yet clear. Nymphs older and younger than stillborn nymphs generally lived to adulthood and were able to reproduce at rates consistent with the offspring of control aphids (Supplementary Fig. 6). Superficially this indicates that the DNA methyltransferases are not required specifically for germ maintenance. However, this may also reflect a period in development in which embryos are refractory to 5-azacytidine treatment or that embryos that were severely affected were resorbed or stillborn. Further studies using RNAi or CRISPR/Cas9 are required to fully define the roles of these two paralogs and dnmt1 in pea aphid oogenesis and embryogenesis.

The co-expression of both *dnmt3* paralogs in the developing germ line raises the question of why both paralogs are maintained in the genome. It may be that *dnmt3x* has been released from the selective pressure towards its ancestral function and has acquired a new function through neofunctionalization. Or, given that both are being expressed in the same way, it may represent a case of subfunctionalization. While *Ap-dnmt3a* has a Cyt C5 DNA methylase domaine (Fig. S4) and is presumed to be functional as a DNA methyltransferase, *Ap-dnmt3x* has a PWWP motif essential for protein-protein interactions and interactions with chromatin (Kucharski et al. 2023a) and a ADDz_Dnmt3 domain (Fig. S4) but this domain is lacking some of the conserved amino acid residues thought to be required for activity as a DNA methyltransferase (Walsh et al. 2010). Therefore, *dnmt3x* may function similarly to the vertebrate Dnmt3L (Walsh et al. 2010). Dnmt3L, while not able to actively methylate DNA itself, stimulates *de novo* methylation through methyltransferases that can (Chédin et al. 2002). As *Ap-dnmt3x* mRNA colocalises with *Ap-dnmt3a*, it is possible that it may play a similar role, interacting with and modulating the activity of *dnmt3a* in establishing methylation patterns. Interestingly, the unusual arrangement of dnmt3s observed within the Aphidomorpha is reminiscent of the system described in *Daphnia magna*. *D. magna* also possess two *dnmt3* orthologs, one of which possesses a putatively functional methyltransferase domain but is lacking a PWWP domain, and the other appears to have a diverged methyltransferase domain, presumed to be non-functional (Lindeman et al. 2019; Nguyen et al. 2020). Given that *D. magna* also exhibit cyclical parthenogenesis (Young 1979; Kleiven et al. 1992) it potentially interesting that the unusual *dnmt3* configurations of aphids and *D. magna* are superficially consistent.

Cyclical parthenogenesis is thought to have evolved in Aphidomorpha over ca. 200 MYA, before the divergence of the *Aphididae*, *Adelgidae* and *Phylloxeridae*, while viviparity in the asexual phase arose in the *Aphididae* specific lineage ca. 140-200 MYA, after the divergence (*Adelgidae* and *Phylloxeridae* reproduce exclusively via oviparous cyclical parthenogenesis) (Davis 2012). As the duplication of Dnmt3 occurred in an ancestor of these three groups it is tempting to speculate that one or both of these paralogs may have a role in mediating parthenogenesis. That the mRNA of both *Ap-dnmt3a* and *Ap-dnmt3x* is detected in the oocytes and germ cells of parthenogenetic viviparous pea aphids and that inhibition of the DNA methyltransferases causes defects in oogenesis and early embryogenesis is consistent with this hypothesis. More work focussing on the role of DNA methyltransferases in oviparous reproduction in aphids, and in other Aphidomorpha will be required to fully test this hypothesis.

## Author Contributions

**KJY:** Conceptualisation, Investigation, Formal Analysis, Visualisation, Writing – Original draft preparation, **SW:** Investigation, Writing – Reviewing and Editing. **EJD:** Conceptualisation, Funding Acquisition, Investigation, Visualisation, Writing – Original draft preparation.

## Acknowledgements

This work was supported by a Royal Society Research Grant (RG170520) to EJD. KJY is supported by a BBSRC White Rose Mechanistic Biology DTP Scholarship (2272108). The authors thank Matthew Chambers and Dr Christopher Cunningham for critical comments on this manuscript and Professor Peter Dearden for critical discussion of the 5’ azacytidine phenotypes.

## Notes

### Competing Interest Statement

The authors have declared no competing interest.

