## Supplementary Figures and Tables for "DNA methylation machinery is involved in development and reproduction in the viviparous pea aphid (*Acyrthosiphon pisum*)"

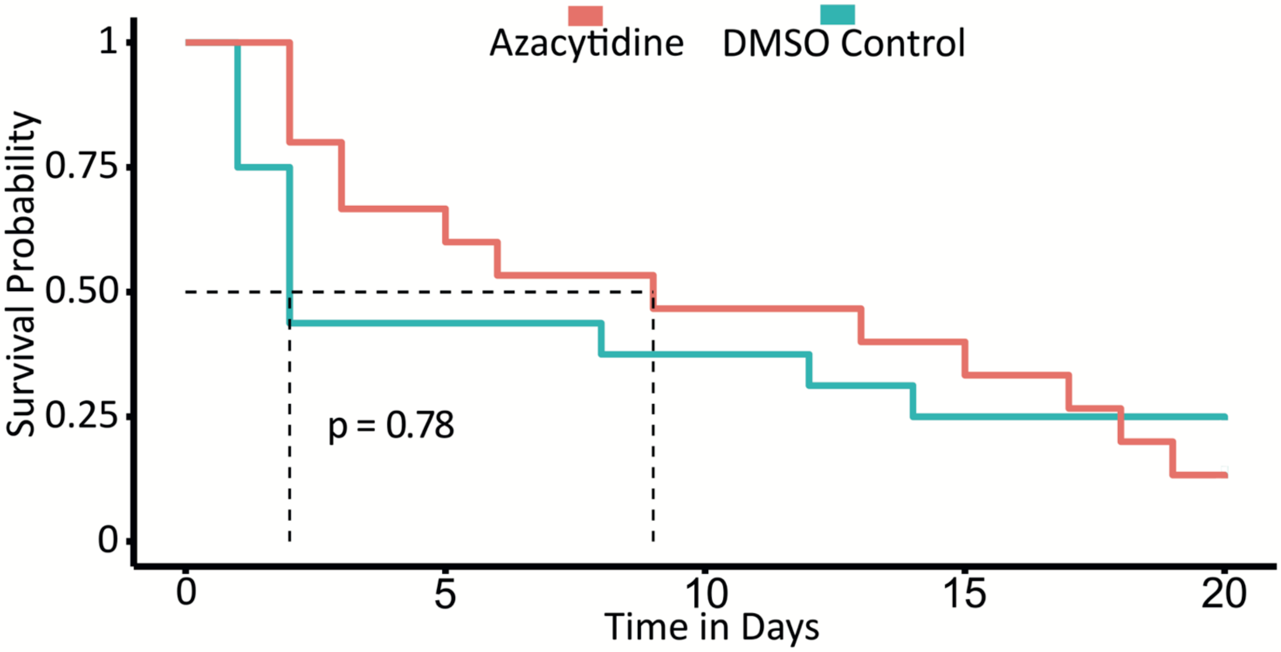


**Supplementary Fig. 1: Kaplan-Meier survival curve of aphids following 5-azacytidine treatment**. There was no significant difference in survival between 5-azacytidine treated aphids and DMSO controls (Cox proportional hazard, p = 0.78).

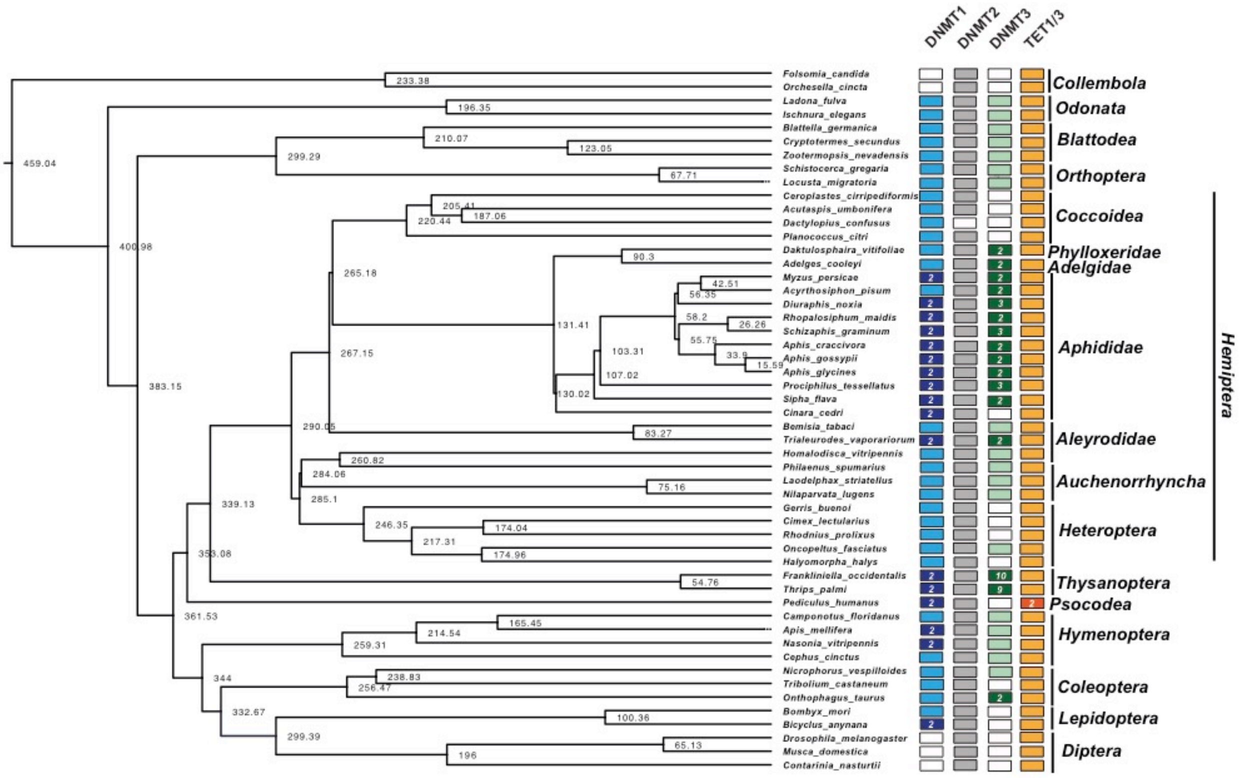


**Supplementary Fig. S2: Conservation of genes involved in DNA methylation across Hemiptera and Thysanoptera.** The phylogenetic relationships between species and divergence times (shown at nodes) are based on TimeTree of Life (Kumar et al. 2022). Lighter colours indicate the presence of a single homolog in the genome, while darker colours indicate the presence of paralogs (the number of paralogs is indicated.  Dnmt1,

Dnmt2 and Tet1 are generally very well conserved across the Hemiptera and Thysanoptera although duplicates of Dnmt1 are often seen, which may indicate the potential for neo or subfunctionilisation of these paralogs. In contrast, dnmt2 and Tet1 are never duplicated.  Dnmt3 exhibits more variability, with independent duplications in Aphidomorpha, Thysanoptera, and a potential species-specific duplication in *Trialeurodes vaporariorum*, along with losses in the Coccoidea and multiple losses in the Heteroptera

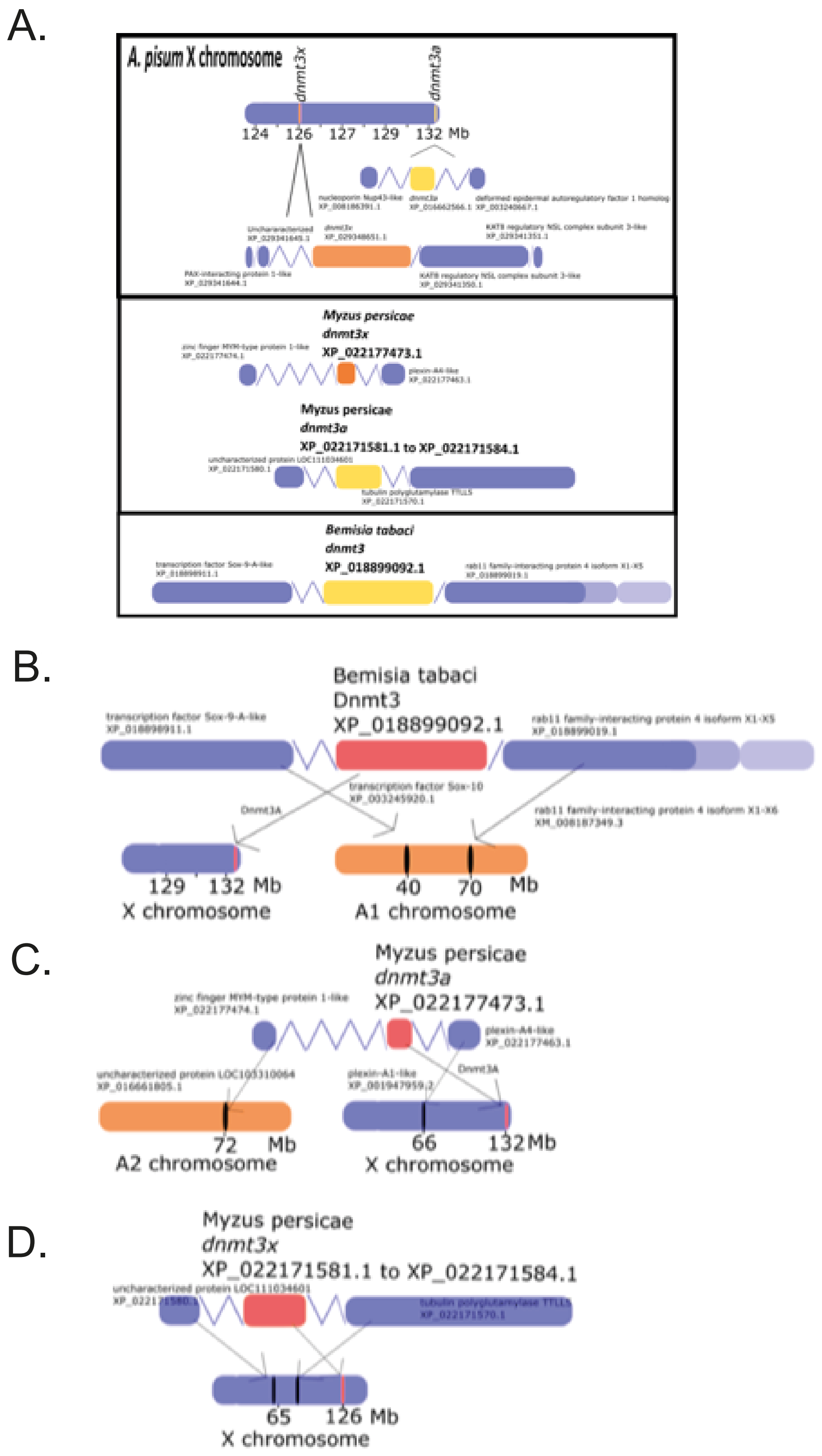


**Supplementary Fig. S3:** Schematic representation of synteny based on *A. pisum* dnmt3a and dnmt3x and flanking genes, with mapping of orthologues of *A. pisum* Dnmt3x and/or

Dnmt3a endoing genes and the genes directly flanking them (identified using NCBI genome browser) from *B. tabaci*, a non-aphid hemipteran bug, and *M. persicae* (another aphid) to *A. pisum* chromosomes (using a chromosome-level *A. pisum* genome assembly). *B. tabaci* possesses only a single Dnmt3, more homologous to *A. pisum* Dnmt3a, and the genes flanking it do not appear on the *A. pisum* X chromosome (where dnmt3a and dnmt3x are located). For *M. persicae* flanking genes, three out of four appear on the X chromosome, nearby to dnmt3a and dnmt3x, but one appears on the A2 chromosome.

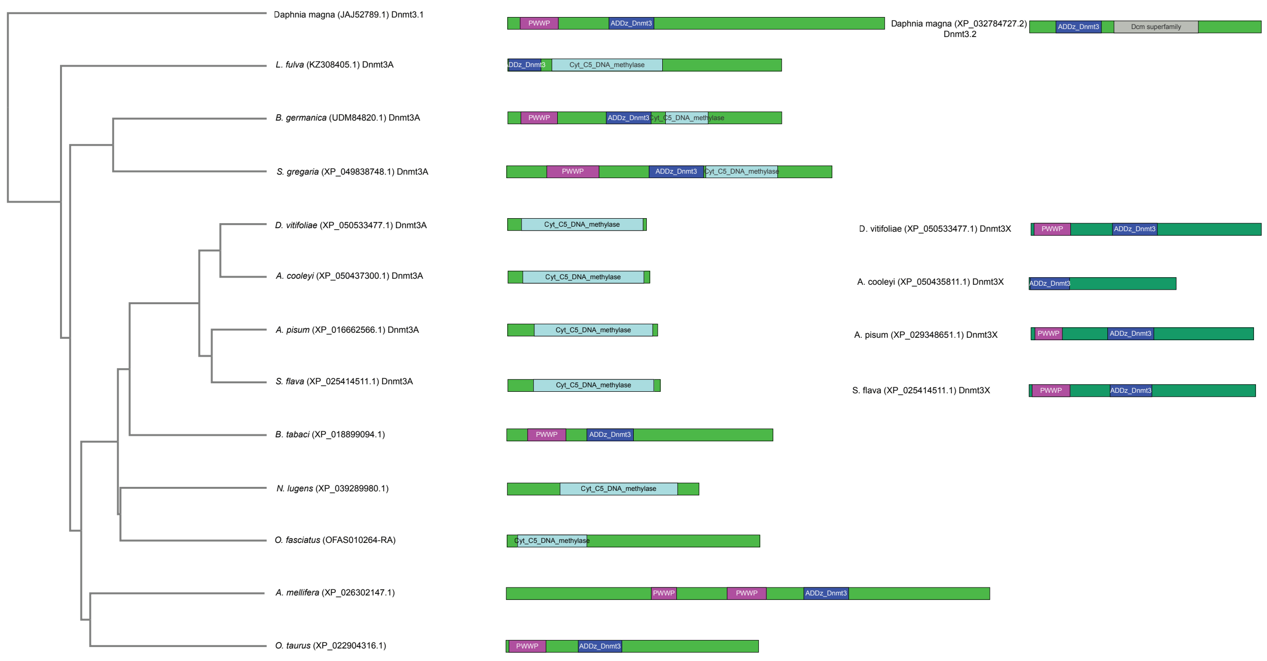


**Supplementary Fig. S4: Domain structure of insect Dnmt3 proteins superimposed on species phylogeny.** Domains identified by Batch CD-search at NCBI, and visualised using IBS 2.0 (Xie et al. 2022)

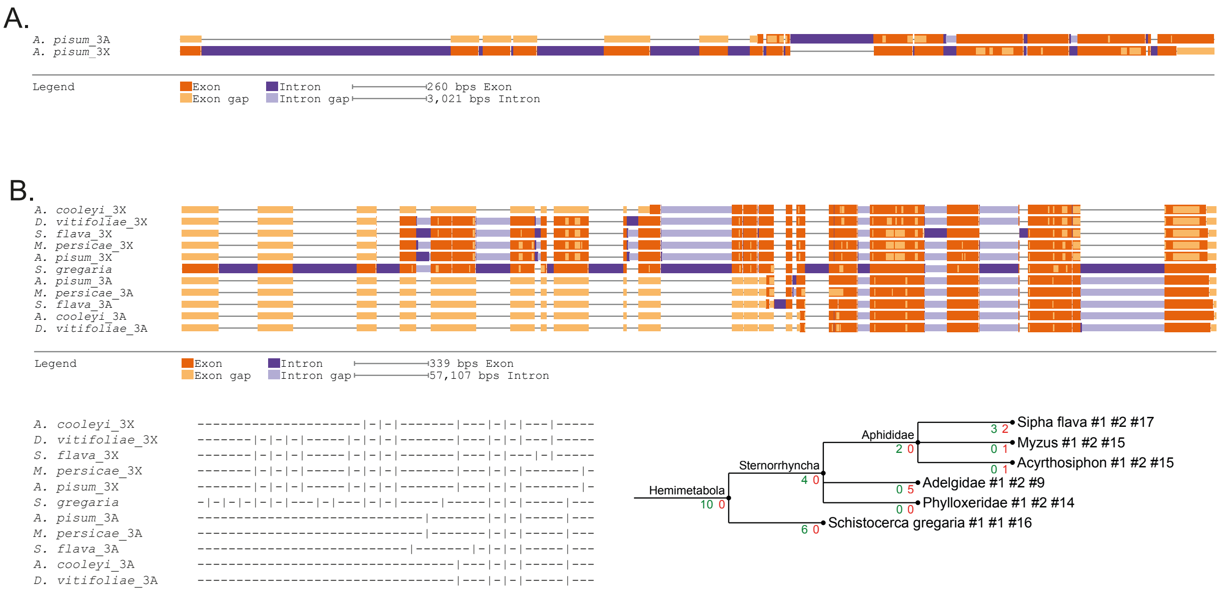


**Supplementary Fig. S5: Conservation of Intron/Exon structure.** A) comparison of intron/exon structure between the pea aphid paralogs of dnmt3. B) comparison of the intron/exon structure between three aphid species (*S. flava, M. persicae and A. pisum*) with representatives of the phylloxera (*D.vitifoliae*) and adelgids *(A. cooleyi*). Gene structures were generated using https://www.webscipio.org/search and then I were compared using https://genepainter.motorprotein.de/genepainter.

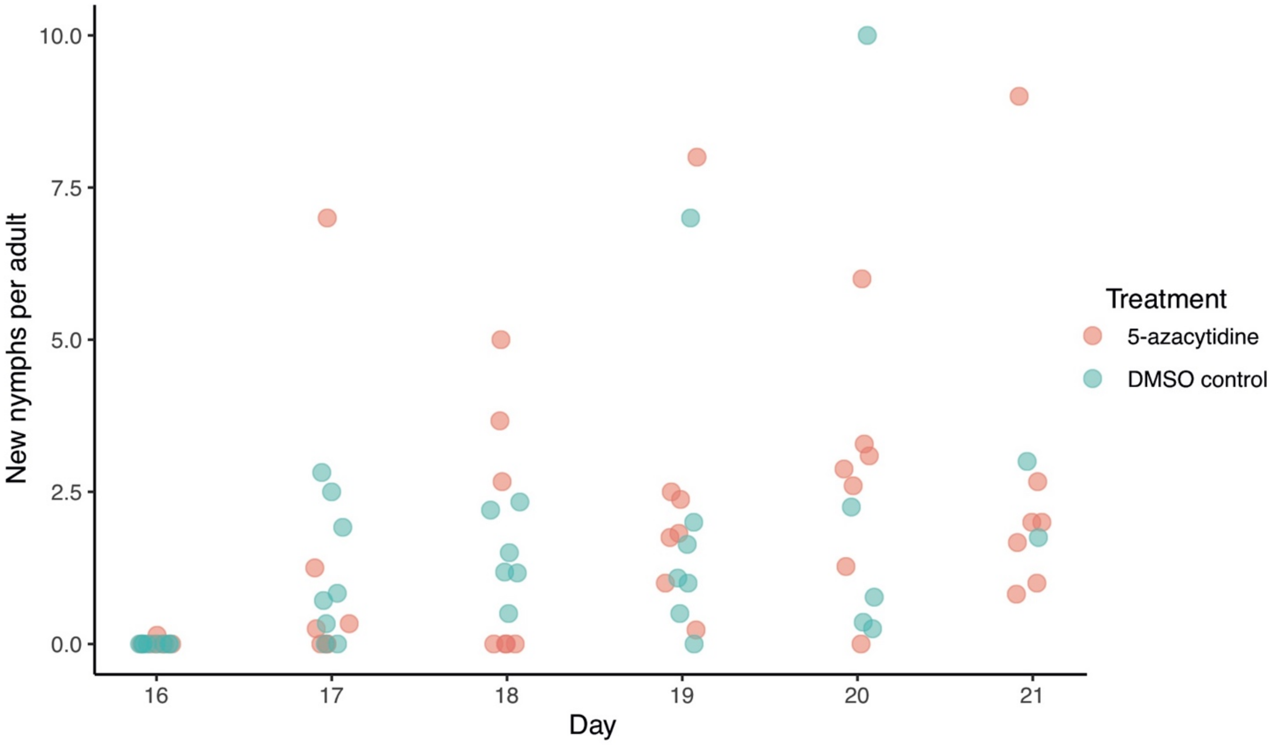


**Supplementary Fig. 6: Fecundity of nymphs produced by 5-azacytidine treated and DMSO control mothers.**

**Supplementary Table 1:** Preparation of aphid artificial diet (Prosser and Douglas 1992). Aphid artificial diet was made by first combining 5 ml of amino acid solution with 0.1 ml of mineral solution (11 mg $FeCl_3.6H_2O$, 2 mg $CuCl_2.4H_2O$, 4 mg $MnCl_2.6H_2O$, and 17 mg $ZnSO_4$ dissolved in 10 ml dd$H_2O$), and 0.5 ml of vitamin solution (0.1 mg biotin, 5 mg pantothenate, 2 mg folic acid, 10 mg nicotinic acid, 2.5 mg pyridoxine, 2.5 mg thiamine, 50 mg choline, and 50 mg myo-inositol dissolved in 5 ml ddH20). Then adding 3 ml sucrose stock solution (10 mg ascorbic acid, 1 mg citric acid, 1.7 g sucrose, and 20 mg $MgSO_4.7H_2O$ dissolved in 3 ml total volume dd$H_2O$), followed by 115 mg $K_2PO_4$ dissolved in 1 ml dd$H_2O$, and then made up to 10 ml with dd$H_2O$. The pH was confirmed as being between 7.0 and 7.5. The solution was then filter-sterilised into sterile plastic tubes and stored at – 20 °C. Amino acid stock solution composition

| **Amino Acid** | | | **mg amino acid to give final amino acid [illegible]** |
|---|---|---|---|
| Alanine | ALA | A | 50.8 |
| Asparagine | ASN | N | 213.9 |

| | | | |
|---|---|---|---|
| Aspartate/Aspartic Acid | ASP | D | 189.7 |
| Cysteine | CYS | C | 42.5 |
| Glutamic Acid | GLU | E | 123.6 |
| Glutamine | GLN | Q | 241.1 |
| Glycine | GLY | G | 9.0 |
| Proline | PRO | P | 65.6 |
| Serine | SER | S | 59.9 |
| Tyrosine | TYR | Y | 10.9 |
| Arginine | ARG | R | 300.2 |
| Histidine | HIS | H | 182.4 |
| Isoleucine | ISO | I | 114.1 |
| Leucine | LEU | L | 114.1 |
| Lysine | LYS | K | 158.9 |
| Methionine | MET | M | 42.5 |
| Phenylalanine | PHE | F | 47.1 |
| Threonine | THR | T | 103.6 |
| Tryptophan | TRP | W | 58.2 |
| Valine | VAL | V | 101.9 |
